## Supporting Information for "The CLIP-domain serine protease CLIPC9 regulates melanization downstream of SPCLIP1, CLIPA8, and CLIPA28 in the malaria vector *Anopheles gambiae*"

Gregory L. Sousa *et al.*

**<u>Figures and Legends</u>**

S1 Figure

S2 Figure

S3 Figure

S4 Figure

S5 Figure

S6 Figure

S7 Figure

S8 Figure

S9 Figure

**<u>Tables</u>**

S1 Table

S2 Table

S3 Table

**<u>Raw Data</u>**

S1 Appendix

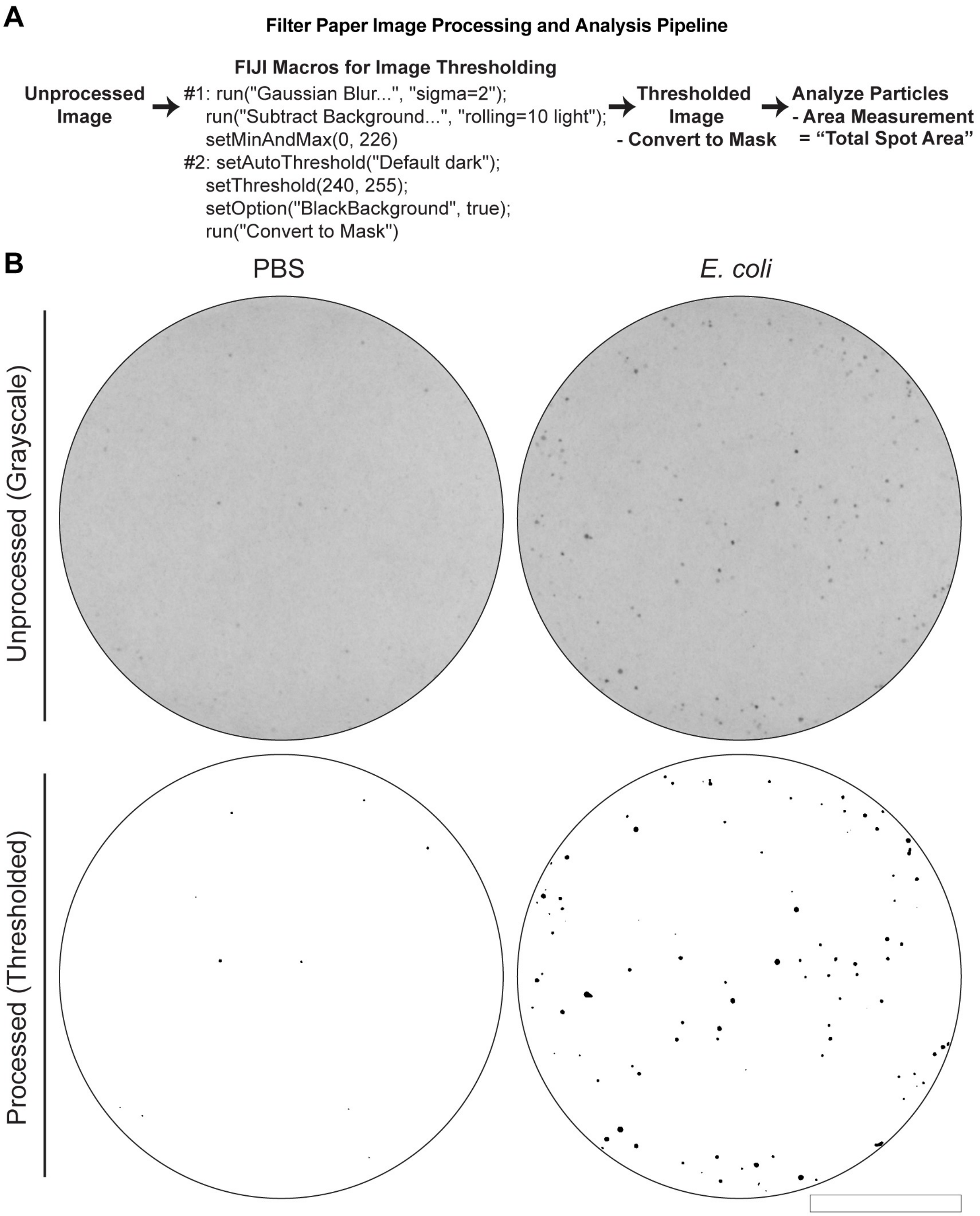


**S1 Figure. Filter paper image processing workflow. (A)** Filter paper image processing and analysis pipeline to generate and quantify thresholded images in FIJI[83]. **(B)** Unprocessed (top row, grayscale) and processed (bottom row, thresholded) filter papers from 50 mosquitoes 12 h after PBS or *E. coli* injection. Scale bar is 2.5 cm.

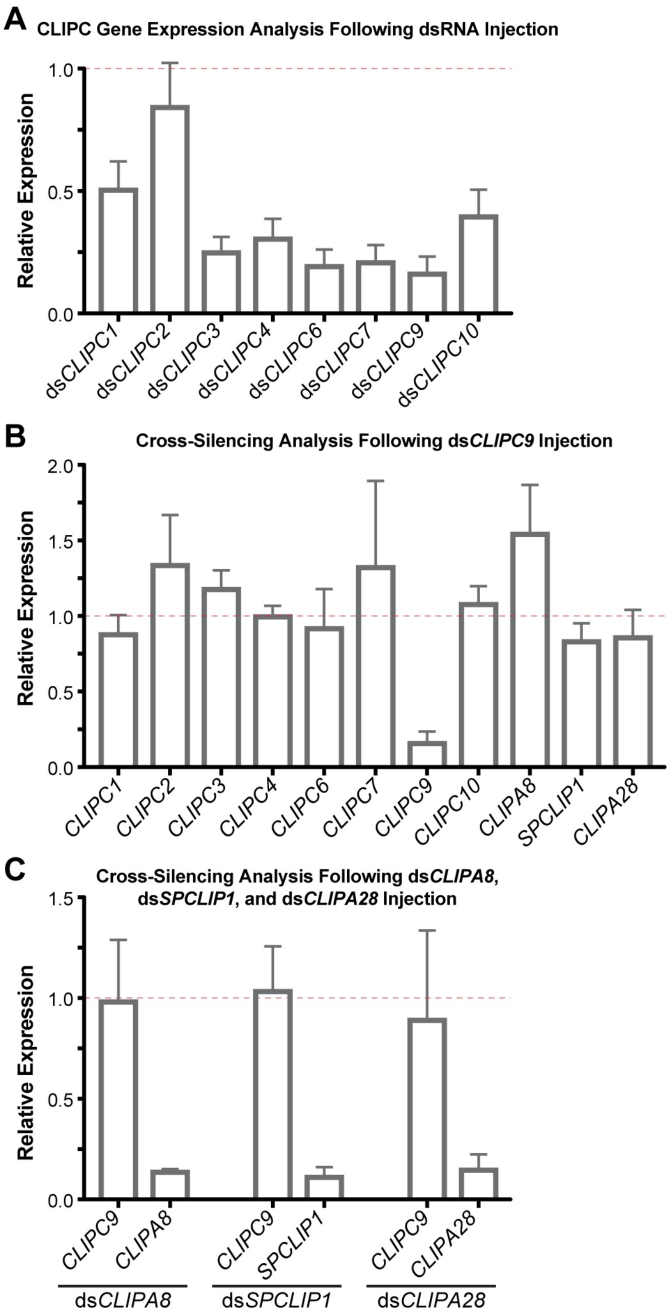


**S2 Figure. Experimental gene knockdown analyses. (A)** CLIPC screen gene expression analysis 3-4 d after dsRNA administration. **(B)** Cross-silencing analysis of the CLIPC candidates and the CLIP-SPH pathway members following ds*CLIPC9* treatment. CLIPC9 gene expression was included as a positive control. **(C)** Cross-silencing analysis of CLIPC9 following ds*CLIPA8*, ds*SPCLIP1*, and ds*CLIPA28* treatments. Expression values for CLIPA8, SPCLIP1, and CLIPA28 were included as positive controls. All mean qRT-PCR expression values are relative to their respective ds*LacZ* treatment groups, which are set to 1.0 and denoted by the red dotted line. All error bars are ± SD. All data are pooled from three independent biological replicates.

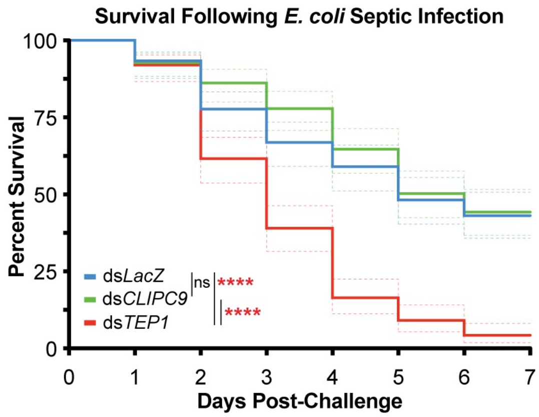


**S3 Figure. CLIPC9 silencing does not affect survival following *E. coli* septic infection.** Mosquito survival was tracked for 7 d following septic infection with *E. coli* at OD 3.2 in ds*LacZ*-, ds*CLIPC9*-, and ds*TEP1*-treated mosquitoes. TEP1 knockdowns were included as a positive control for reduced survival following *E. coli* infection[48]. Experimental groups were compared to the ds*LacZ* group with the Log-rank test. Asterisks denote statistical significance (****p ≤ 0.0001). Curves are averages from three independent biological replicates and dotted lines are 95% confidence intervals.

**A**

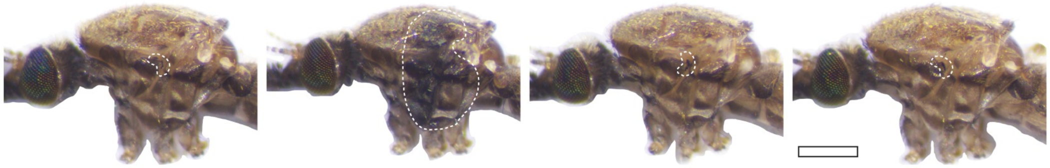


| | Enhanced Thoracic Melanization | | | | |
|---|---|---|---|---|---|
| Group | No | Yes | n | Total (%) | p-value |
| ds*LacZ* | 194 | 0 | 194 | 0.0 | |
| ds*CTL4* | 127 | 62 | 189 | 32.8 | <0.0001 |
| ds*CLIPC9* | 193 | 0 | 193 | 0.0 | ns |
| ds*CLIPA8* | 194 | 0 | 194 | 0.0 | ns |

**B**

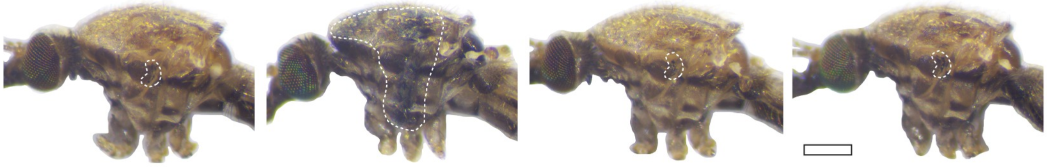


| | Enhanced Thoracic Melanization | | | | |
|---|---|---|---|---|---|
| Group | No | Yes | n | Total (%) | p-value |
| ds*LacZ*/ds*LacZ* | 189 | 0 | 189 | 0.0 | nd |
| ds*LacZ*/ds*CTL4* | 93 | 86 | 179 | 48.0 | |
| ds*LacZ*/ds*CLIPC9* | 171 | 15 | 186 | 8.1 | <0.0001 |
| ds*CLIPA8*/ds*CTL4* | 170 | 8 | 178 | 4.5 | <0.0001 |

**S4 Figure. Injection with ds*CTL4* causes an enhanced injection site melanization response that requires CLIPC9 and CLIPA8.** Brightfield photomicrographs of mosquito thoraxes 3-4 d after **(A)** single or **(B)** double gene knockdown. Single gene knockdown data were compared to ds*LacZ* control using Fisher's exact test. The ds*LacZ*/ds*LacZ* treatment group was excluded from the double gene knockdown analysis. The ds*CLIPC9*/ds*CTL4* and ds*CLIPA8*/ds*CTL4* treatment groups were each compared to the ds*LacZ*/ds*CTL4* group using Fisher's exact test. The regions of injection-induced melanization are outlined in white. Scale bar is 500 µm in **(A)** and 200 µm in **(B).** All data are compiled from three independent biological replicates. Abbreviations: ns = not significant; nd = not determined.

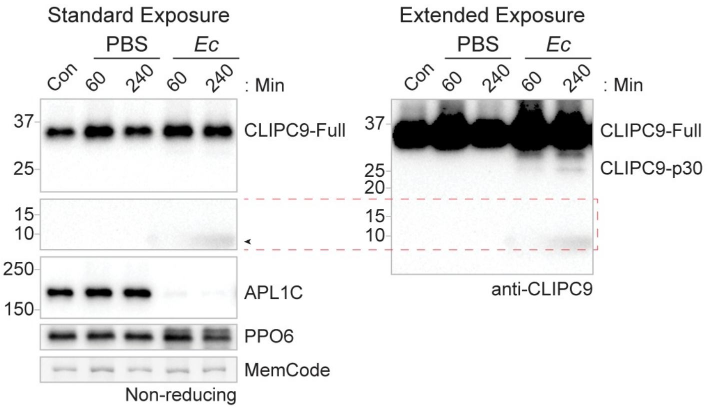


**S5 Figure. CLIPC9 cleavage products are disulfide linked.** The septic infection-induced cleavage of CLIPC9 was assessed by non-reducing western analysis of naïve/control hemolymph (Con) and hemolymph 60 and 240 min after PBS or *E. coli* (*Ec*) challenge. Blots were probed with antibodies against APL1C and PPO6 to confirm *E. coli* exposure and equal protein loading, respectively. Black arrowhead denotes a non-specific haze associated with *E. coli* injection in western analysis. Membranes were MemCode stained as an additional loading control. Image is representative of three independent biological replicates.

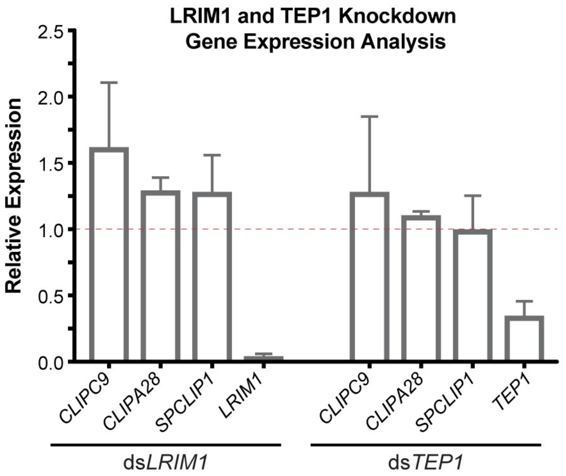


**S6 Figure. Gene expression analysis in LRIM1 and TEP1 knockdowns.** Mean qRT-PCR expression analysis of CLIPC9, CLIPA28, and SPCLIP1 3-4 d after ds*LRIM1* and ds*TEP1* administration. Expression values for LRIM1 and TEP1 were included as positive controls. Expression values are relative to the ds*LacZ* treatment group, which is denoted by the red dotted line set to 1.0. Error bars are ± SD. Experimental replicates are from three independent mosquito generations.

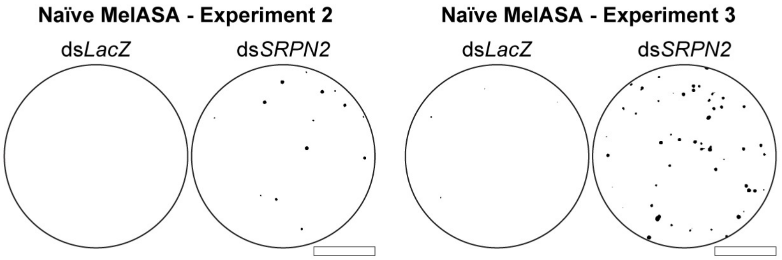


**S7 Figure. Enhanced MelASA spot generation in ds*SRPN2*-treated mosquitoes.** Additional naïve Mini-MelASA replicates 4 d after gene knockdown in ds*SRPN2*- and ds*LacZ*-treated mosquitoes. Filter papers were removed after 12 h. Scale bar is 1.3 cm.

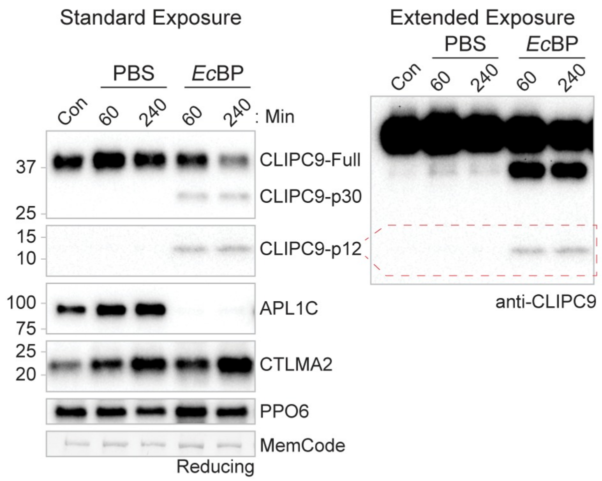


**S8 Figure. CLIPC9 is cleaved after *E. coli* bioparticle injection.** Challenge-induced cleavage of CLIPC9 was assessed by reducing western analysis on whole hemolymph 60 and 240 min after PBS or *E. coli* bioparticle injection (*Ec*BP) and from naïve/control (Con) mosquitoes. Blots were probed against APL1C, CTLMA2, and PPO6 to confirm the *Ec*BP challenge, adequate sample reduction, and equal protein loading, respectively. Membranes were MemCode stained as an additional loading control. Western image is representative of experiments performed on two independent biological replicates.

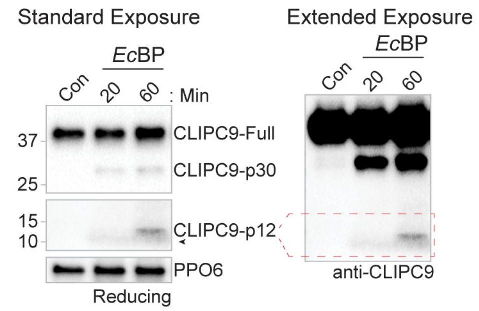


**S9 Figure. CLIPC9 p12 fragment is detectable 60 min after *E. coli* bioparticle injection.** Challenge-induced cleavage of CLIPC9 was assessed by reducing western analysis on whole hemolymph 20 and 60 min after *E. coli* bioparticle (*Ec*BP) injection and from naïve/control (Con) mosquitoes. Blots were probed against PPO6 to confirm equal protein loading. Western image is representative of four independent biological replicates.

**S1 Table. T7 primers used for dsRNA template synthesis**

| AGAP ID | Name | Sequence |
|---|---|---|
| n/a | LacZ T7 F | taatacgactcactatagggAGAATCCGACGGGTTGTTACT |
| --- | LacZ T7 R | taatacgactcactatagggCACCACGCTCATCGATAATTT |
| AGAP010731 | CLIPA8 T7 F | taatacgactcactatagggGCAGAACGATGGCTCAGATG |
| --- | CLIPA8 T7 R | taatacgactcactatagggTGCTCGTTGGACGAGTAGAA |
| AGAP010730 | CLIPA28 T7 F | taatacgactcactatagggGAATGGGACATCAGCACCAC |
| --- | CLIPA28 T7 R | taatacgactcactatagggCGCAGGGTGTAGAACGGT |
| AGAP028725 | SPCLIP1 T7 F | taatacgactcactatagggGTCACCGAACACGGCCAAC |
| --- | SPCLIP1 T7 R | taatacgactcactatagggATCGAAGCTGATCGGATCGGG |
| AGAP005335 | CTL4 T7 F | taatacgactcactatagggTGGTTTGATGCCGTGTCCT |
| --- | CTL4 T7 R | taatacgactcactatagggAATAAATTGTCTCGGTTCATCATC |
| AGAP006348 | LRIM1 T7 F | taatacgactcactatagggAATATCTATCTCGCGAACAATAA |
| --- | LRIM1 T7 R | taatacgactcactatagggTGGCACGGTACACTCTTCC |
| AGAP010815 | TEP1 T7 F | taatacgactcactatagggTTTGTGGGCCTTAAAGCGCTG |
| --- | TEP1 T7 R | taatacgactcactatagggACCACGTAACCGCTCGGTAAG |
| AGAP006911 | SRPN2 T7 F | taatacgactcactatagggCGAGGGCGCGGTCATTACG |
| --- | SRPN2 T7 R | taatacgactcactatagggCAGCATTGTTCCGAGGGTTTCATC |
| AGAP008835 | CLIPC1 T7 F | taatacgactcactatagggGAGTATGGGCAGGCTGTGTT |
| --- | CLIPC1 T7 R | taatacgactcactatagggAAGTTTGTGGAAGTTAGGCAGTG |
| AGAP004317 | CLIPC2 T7 F | taatacgactcactatagggGAGACTACCAGAGCGCATCC |
| --- | CLIPC2 T7 R | taatacgactcactatagggGTCGCCAAACTCGTCGTT |
| AGAP004318 | CLIPC3 T7 F | taatacgactcactatagggGCCGACGATCGATAGAAACT |
| --- | CLIPC3 T7 R | taatacgactcactatagggGCAGCGCAAAGAGATCAGAC |
| AGAP000573 | CLIPC4 T7 F | taatacgactcactatagggGAGTTCAGCGGCAACGAT |
| --- | CLIPC4 T7 R | taatacgactcactatagggTGCACCTGCTTGATCTCGTA |
| AGAP000571 | CLIPC5 T7 F | taatacgactcactatagggACGTACGATGGGACGCAG |
| --- | CLIPC5 T7 R | taatacgactcactatagggAGCAGGAAGCGAACCGTTAT |
| AGAP000315 | CLIPC6 T7 F | taatacgactcactatagggCAGACGCTTTACGAGGGAGA |
| --- | CLIPC6 T7 R | taatacgactcactatagggTACTTGGGCACCTGTTCGC |
| AGAP003689 | CLIPC7 T7 F | taatacgactcactatagggCAATTACGGTACAGCTGGGAA |
| --- | CLIPC7 T7 R | taatacgactcactatagggCCTCCGTTAGCGTAAGGTTG |
| AGAP004719 | CLIPC9 T7 F | taatacgactcactatagggGGTGCAGTAAGAAGGCCCAT |
| --- | CLIPC9 T7 R | taatacgactcactatagggACTGCATGTCCAAGCAATCC |
| AGAP000572 | CLIPC10 T7 F | taatacgactcactatagggCTCATCTCGTCCCGGTTTCT |
| --- | CLIPC10 T7 R | taatacgactcactatagggGCGATATCGTTCTGGTACGTG |

**S2 Table. Primers used for qRT-PCR analysis**

| AGAP ID | Name | Sequence |
|---|---|---|
| AGAP010592 | S7 qPCR F | GTGCGCGAGTTGGAGAAGA |
| --- | S7 qPCR R | ATCGGTTTGGGCAGAATGC |
| AGAP010731 | CLIPA8 qPCR F | GATCGATTCGACGACCAACT |
| --- | CLIPA8 qPCR R | GCAGGTCGACTCGCTTTAAC |
| AGAP010730 | CLIPA28 qPCR F | ATAAAGCATGCCCAAACCAC |
| --- | CLIPA28 qPCR R | CTGACAGCACACCAGCAGAT |
| AGAP028725 | SPCLIP1 qPCR F | CTTGCTGAACGACGACGATA |
| --- | SPCLIP1 qPCR R | TCCTGTTCCTCCTCGTCACT |
| AGAP005335 | CTL4 qPCR F | GCACGGGTACAGGGCTACTA |
| --- | CTL4 qPCR R | GCGTGGTGTACAGCTTTCCT |
| AGAP006348 | LRIM1 qPCR F | CATCCGCGATTGGGATATGT |
| --- | LRIM1 qPCR R | CTTCTTGAGCCGTGCATTTTC |
| AGAP010815 | TEP1 qPCR F | AAAGCTGTTGCGTCAGGG |
| --- | TEP1 qPCR R | TTCTCCCACACACCAAACGAA |
| AGAP006911 | SRPN2 qPCR F | CCATACGCGCAGCTACTACA |
| --- | SRPN2 qPCR R | ACCTCGATGAAGTCGTCCAC |
| AGAP008835 | CLIPC1 qPCR F | CACCCCGAGTACAAGCAAAC |
| --- | CLIPC1 qPCR R | CGTGCGTAGGGTGAGAAGAT |
| AGAP004317 | CLIPC2 qPCR F | CTCGTTCGGTATTGGCTGTG |
| --- | CLIPC2 qPCR R | TGCCCTCGATCCAGTCTATG |
| AGAP004318 | CLIPC3 qPCR F | CCCATCGGAAGTGCTAAACG |
| --- | CLIPC3 qPCR R | TGGCACCAAGATACTCCGTT |
| AGAP000573 | CLIPC4 qPCR F | ATTGGACCGTCGATCAACCT |
| --- | CLIPC4 qPCR R | GTGCCCTTCATCAGCTTGTC |
| AGAP000315 | CLIPC6 qPCR F | AAGGGTACGGCATCATCGAT |
| --- | CLIPC6 qPCR R | TGGTTACACTTTTGCCACGG |
| AGAP003689 | CLIPC7 qPCR F | CTATTGTCACGTTGCGCTCA |
| --- | CLIPC7 qPCR R | GATTTCCAGTCTTCCGGCAC |
| AGAP004719 | CLIPC9 qPCR F | ATTGCAATCCGGTGAACATT |
| --- | CLIPC9 qPCR R | TGGGTTCGAGATGAAGCTCT |
| AGAP000572 | CLIPC10 qPCR F | GAGCAGAACGAAACGGTGG |
| --- | CLIPC10 qPCR R | CGTGATGCCGACCAGATAGT |

**S3 Table. Cloning primers used for protein production**

| AGAP ID | Name | Sequence |
|---|---|---|
| AGAP004719 | CLIPC9-FL BamHI F | CATGGATCCGCAGTAAGAAGGCCCATTAATATAC |
| --- | CLIPC9-FL HindIII R | CATAAGCTTTTAGTTATCCCTAGTAGCACTAAC |
| AGAP004719 | CLIPC9-FL NcoI F | GATACCATGGCAGTAAGAAGG |
| --- | CLIPC9-FL XhoI R | CGTTCTCGAGTTAGTTATCCCTAGTAG |
| AGAP010730 | CLIPA28-FL BamHI F | CATGGATCCCAAGACATTGAAGAAGAACTGAGATG |
| --- | CLIPA28-FL HindIII R | CATAAGCTTTTACAATTTTATATCAAAACTCTC |

**S1 Appendix. Counts of viable oocysts and melanized ookinetes in single and double knockdown experiments.**
